# miR-29 is an important driver of aging-related phenotypes

**DOI:** 10.1101/2022.11.29.518429

**Authors:** Vijay Swahari, Ayumi Nakamura, Emilie Hollville, Yu-Han Hung, Matt Kanke, C. Lisa Kurtz, Xurde M. Caravia, Shenghui He, Janakiraman Krishnamurthy, Sahil Kapoor, Varun Prasad, Cornelius Flowers, Matt Beck, Jeanette Baran-Gale, Norman Sharpless, Carlos López-Otín, Praveen Sethupathy, Mohanish Deshmukh

## Abstract

Aging is a consequence of complex molecular changes, but the roles of individual microRNAs (miRNAs) in aging remain unclear. One of the few miRNAs that are upregulated during both normal and premature aging is miR-29. We confirmed this finding in our study in both mouse and monkey models. Follow-up analysis of the transcriptomic changes during normal aging revealed that miR-29 is among the top miRNAs predicted to drive the aging-related gene expression changes. We also showed that partial loss of miR-29 extends the lifespan of *Zmpste24*^*-/-*^ mice, an established model of progeria, which indicates that miR-29 is functionally important in this accelerated aging model. To examine whether miR-29 upregulation alone is sufficient to promote aging-related phenotypes *in vivo*, we generated mice in which miR-29 can be conditionally overexpressed (miR-29TG). We found that miR-29 overexpression in mice is sufficient to drive aging-related phenotypes including alopecia, kyphosis, osteoporosis, senescence, and leads to early lethality. Transcriptomic analysis of both young miR-29TG and old WT mice revealed shared downregulation of genes enriched in extracellular matrix and fatty acid metabolism, and shared upregulation of genes in pathways linked to inflammation. Together, these results highlight the functional importance of miR-29 in controlling a gene expression program that drives aging-related phenotypes.

## Main text

The miR-29 family is known to be induced during normal aging^1-11^. miR-29 was also reported to be elevated in two mouse models of premature aging^1,2^,12 and, interestingly, is also one of only two miRNAs that are reduced in the Ames Dwarf mouse model in which the onset of aging is delayed^13^. Despite this strong correlation of miR-29 levels with normal, premature, and delayed aging, whether miR-29 alone is necessary or sufficient to drive the aging phenotype *in vivo* has not been investigated.

We first sought to characterize the elevation of miR-29 in different models of aging. The miR-29 family is comprised of miR-29a, miR-29b, and miR-29c (expressed from two genomic loci: *miR-29a/b1* and *miR-29b2/c*), and all family members are identical in the seed sequence, which plays an essential role in the targeting of mRNAs^14^. We examined the levels of the miR-29 family in Rhesus macaque liver and found miR-29 levels to increase with age (Figure 1A and Figure supplement 1A). We next measured miR-29 family members in the *Lmna*^*G609G*^ mouse model of progeria^15^ and found that while all three miRNAs are trending upward, only miR-29b is significantly elevated in this model of accelerated aging (Figure supplement 1B). Based on these results, and because miR-29b is expressed from both genomic loci, we focused on miR-29b for our subsequent studies in mice. We showed that miR-29b levels are significantly increased with age in multiple tissues in mice (Figure 1B).

**Figure 1.**
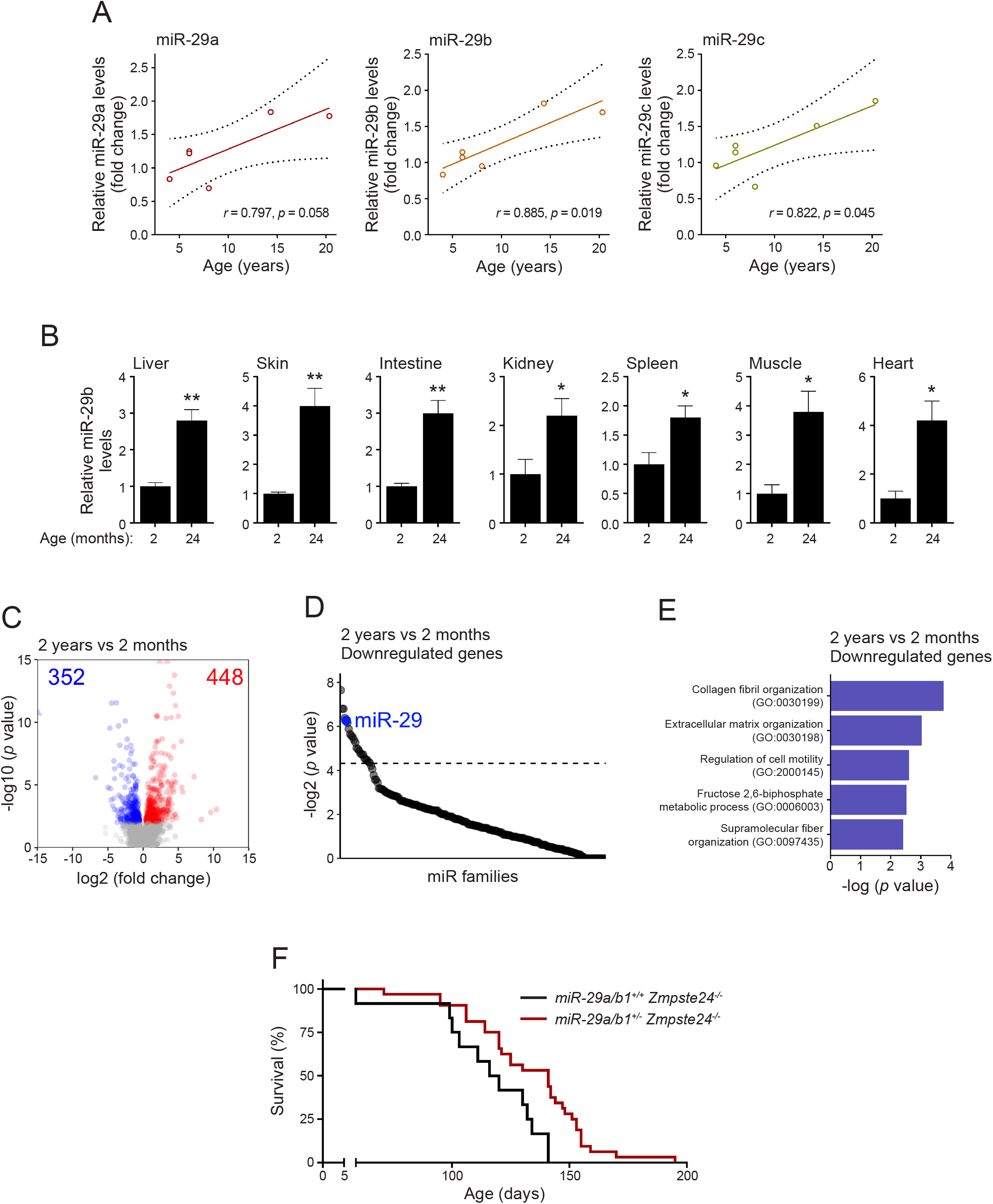
miR-29 is a candidate driver of gene expression changes in aged mice. **(A)** Correlation analysis of the change in levels of miR-29 family members with age in the liver of male Rhesus macaque. Pearson correlation *r* is represented. **(B)** Quantitative PCR analysis for miR-29b in multiple tissues from 2-month and 24-month-old mice. Data shown are mean ± s.e.m. **(C)** Volcano plot showing significantly upregulated (red) and downregulated (blue) genes in liver from old (2 years) mice (n = 5; 3 males, 2 females) compared to liver from young (2 months) mice (n =5; 3 males, 2 females). Filtering criteria: *p* < 0.05, adjusted *p* < 0.2 (Wald test; DESeq2). **(D)** miRHub analysis of the downregulated genes in 2-year-old mouse liver compared to 2-month-old mouse liver. Predicted target sites of the miR-29 family (blue) are significantly enriched in the set of downregulated genes. Dashed line corresponds to *p* = 0.05. **(E)** Pathway enrichment analysis of the genes that are significantly downregulated in old compared to young mouse liver. The analysis was performed based on Gene Ontology (GO) Biological Process 2021. The top 5 processes are represented. **(F)** Kaplan-Meier survival curve of *miR-29a/b1*^*+/+*^ *Zmpste24*^*-/-*^ and *miR-29a/b1*^*+/-*^ *Zmpste24*^*-/-*^ mice (*p* = 0.005, Log-rank/Mantel-Cox test). The median survival is 118.0 days for *miR-29a/b1*^*+/+*^ *Zmpste24*^*-/-*^ and 141.0 days for *miR-29a/b1*^*+/-*^ *Zmpste24*^*-/-*^.

To determine the functional potential of miR-29 to drive molecular changes during aging, we conducted an unbiased query of the genes that are differentially expressed in the liver during normal aging in mice (Figure 1C). We found that the genes significantly downregulated (n = 352, FDR < 0.1) in 2-year-old mice relative to 2-month-old mice are significantly enriched for predicted targets of miR-29 (Figure 1D). Expectedly, the significantly upregulated genes (n = 448, FDR < 0.1) do not harbor any enrichment for miR-29 targets (Figure supplement 2A). We repeated this analysis after stratification by sex and found that miR-29 is the only miRNA with significant enrichment of predicted targets among downregulated genes in both aged males and females (Figure supplements 2C-J). An unbiased pathway analysis of the downregulated genes in aged mice revealed enrichment in collagen and extracellular matrix components, which are well known targets of miR-29^14^ (Figure 1E), while the upregulated genes are enriched in pathways associated with sphingolipid metabolism and neutrophil-mediated immune response (Figure supplements 2B). These results strongly suggest that the gene expression changes during aging could be a consequence of elevated levels and activity of miR-29.

We next examined whether reducing miR-29 levels could extend the survival of the *Zmpste24*^*-/-*^ mice, an established model of progeria^16-20^. The *Zmpste24*^*-/-*^ mouse model is particularly relevant because all three miR-29 family members have previously been shown to be aberrantly elevated in *Zmpste24*^-/-^ mice^1^. Unfortunately, complete deletion of miR-29 (deletion of both *miR-29a/b1* and *miR-29b2/c* loci) results in early lethality^21,22^, precluding us from examining the consequences of a total loss of miR-29 in the context of aging. We, therefore, crossed the *Zmpste24*^*-/-*^ mice with mice deficient for only the *miR-29a/b1* locus to examine the consequence of partial reduction of miR-29 on the survival of the *Zmpste24*^*-/-*^ mice. We used mice deleted for the *miR-29a/b1* locus because it has been reported as the dominant source of miR-29b^23^. Strikingly, we found that the *miR-29a/b1*^*+/-*^ *Zmpste24*^*-/-*^ mice exhibit a significant lifespan extension compared with *miR-29a/b1*^*+/+*^ *Zmpste24*^*-/*-^ mice (median survival 141 *versus* 118 days, p = 0.005), indicating that miR-29 is important for driving premature aging in this progeria model (Figure 1F).

Next, we sought to determine whether increased expression of miR-29 is sufficient to drive aging-related phenotypes *in vivo*. To examine this, we generated transgenic mice that overexpress miR-29b (henceforth referred to as miR-29) ubiquitously under a tetracycline-inducible promoter (miR-29TG) (Figure 2A). Strikingly, by 2 months, overexpression of miR-29 led to an overt aging-like phenotype (Figure 2B). The levels of miR-29 overexpression (2-4 fold) obtained with doxycycline administration in these miR-29TG mice are comparable to the fold increase in miR-29 seen with age in WT mice (Figure 2C). The miR-29TG mice exhibited growth retardation and premature death, with mice dying by 80 days (Figures 2D and 2E).

**Figure 2.**
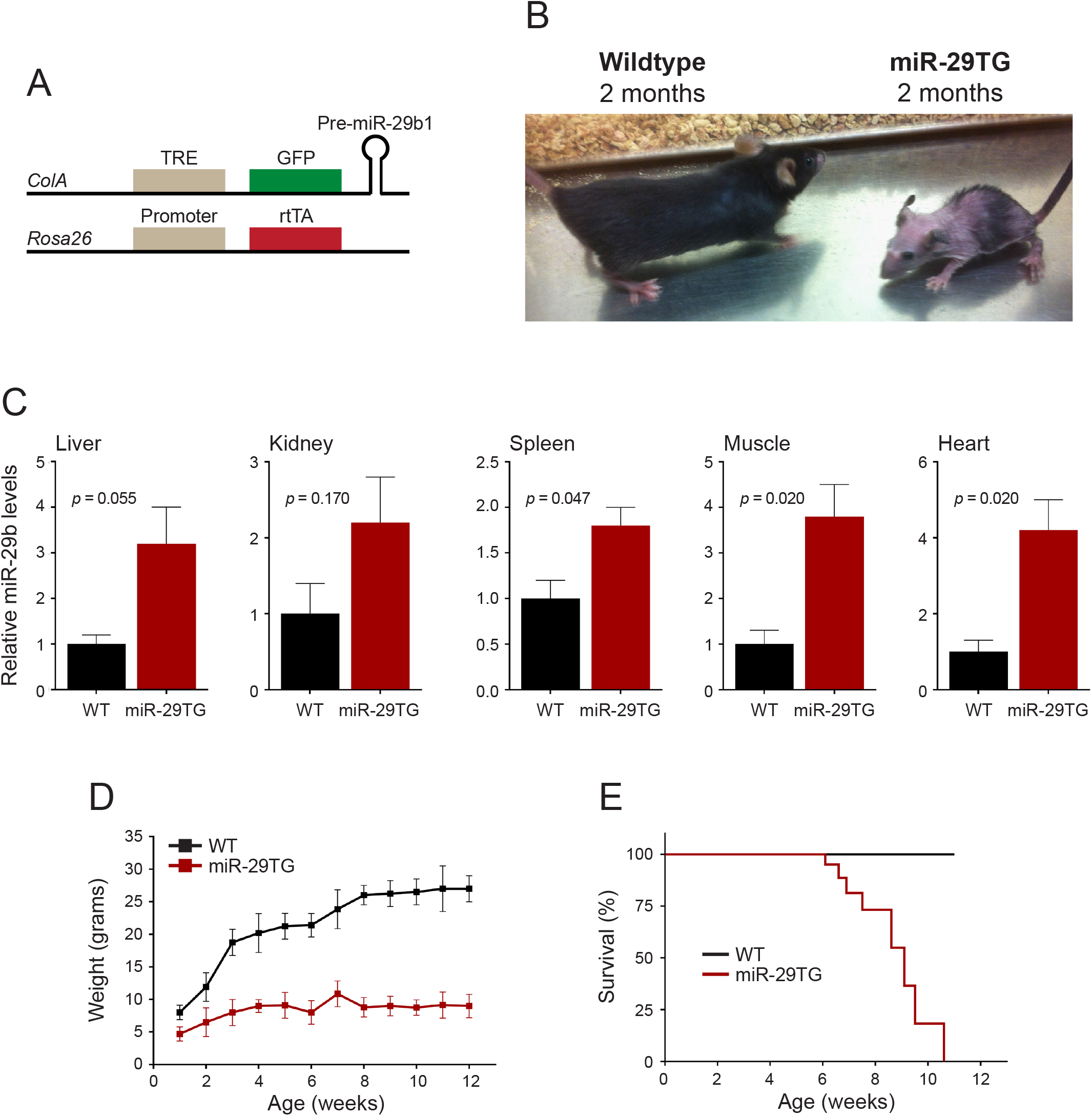
Overexpression of miR-29 results in growth retardation and premature death in mice. **(A)** Constructs used to generate the miR-29 transgenic (miR-29TG) mice. miR-29b and GFP are expressed under the control of tetracycline response element (TRE). In the presence of doxycycline, the tetracycline response transactivator (rtTA) enables TRE-driven expression of miR-29b and GFP. **(B)** Wildtype (left) and miR-29TG (right) mice at 2 months; miR-29 was induced with doxycycline starting at postnatal day 1. **(C)** Quantitative PCR analysis of miR-29b levels in the indicated tissues of miR-29TG mice as compared to WT mice (2 months). Weights **(D)** and Kaplan-Meier survival curve **(E)** of WT and miR-29TG mice. The median survival of miR-29TG mice is 91 days. Data shown are mean ± s.e.m. in **(C)** and **(D)**.

Starting at approximately one month of age, miR-29TG mice began to display premature graying of hair as well as extensive kyphosis that was confirmed using Computed Tomography (CT) imaging analysis (Figures 3A and 3B). The miR-29TG mice also exhibited severe lipodystrophy characterized by little or absent inguinal fat pads at 2 months (Figure 3C). Further, these mice showed reduced skin thickness in all layers of the dermis and epidermis (Figures 3D and 3E). A prominent feature of both normal aging and progeria is the development of osteoporosis. We conducted bone densitometry studies using micro-CT analysis and found that miR-29TG mice indeed have signs of severe osteoporosis as characterized by reduced cortical and trabecular bone volume, reduced cortical bone mineral and total density, and reduced trabecular bone thickness (Figures 3F-I, Figure supplements 3A-C). Similar reductions in bone volume have been observed in the *Lmna*^*G609G*^ mouse model of progeria^15^. In addition, thymus weight and cellularity were reduced in miR-29TG mice, and the normal sequence of thymocyte maturation was disrupted (Figure supplements 3D-F). Interestingly, the frequency of phenotypic hematopoietic stem cells (HSCs) in the bone marrow and spleen was significantly reduced in the miR-29TG mice (Figure supplement 3G). Thus, miR-29 overexpression results in many phenotypes that are known to be associated with aging^24-27^.

**Figure 3.**
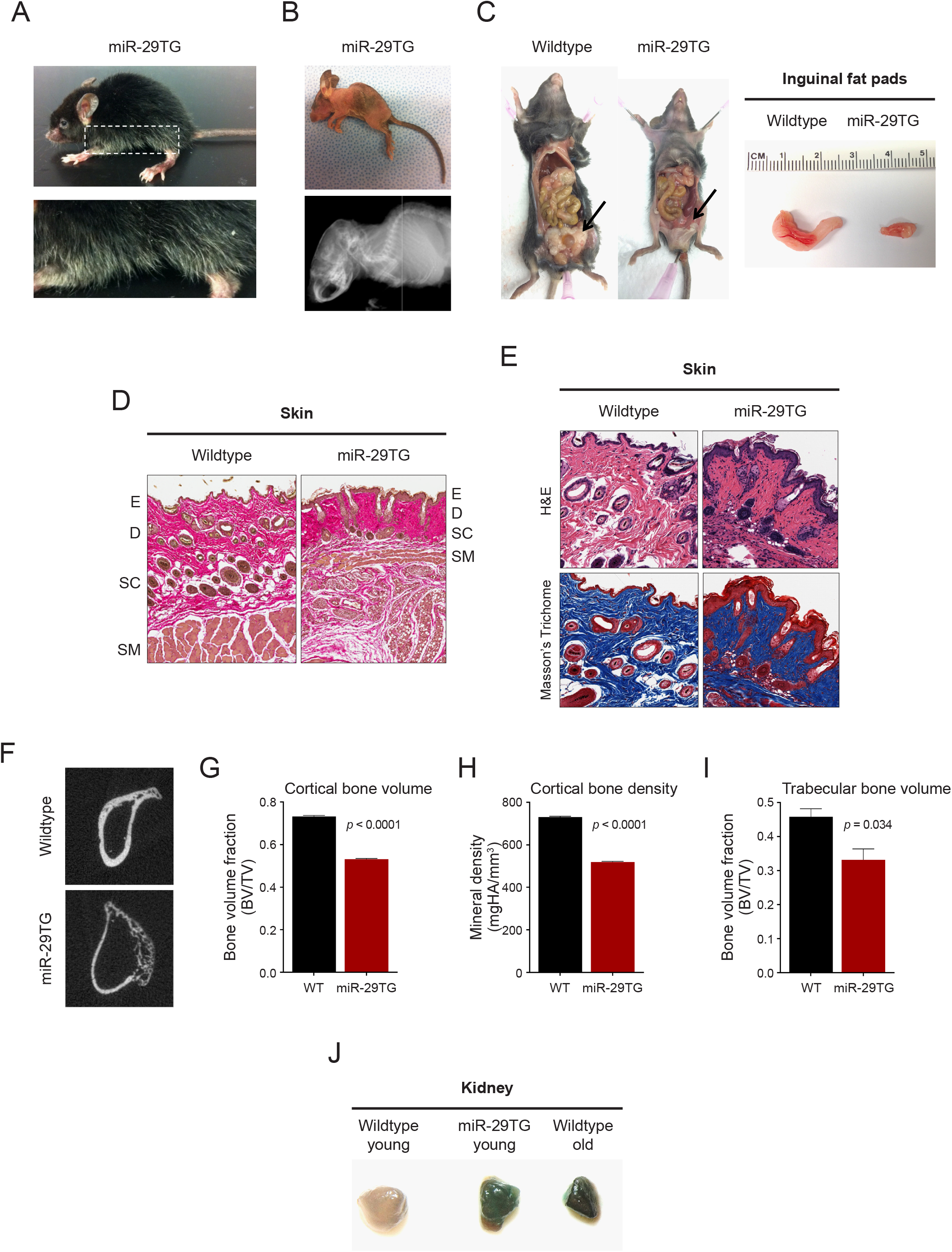
miR-29TG mice exhibit signs of premature aging. **(A)** miR-29TG mice develop hair graying and **(B)** kyphosis (top, photograph; bottom, CT scan) by 2 months. **(C)** Left: lipodystrophy observed in 2-month-old miR-29TG mice (arrows). Right: inguinal fat pads of wildtype and miR-29TG mice. **(D)** Skin of wildtype and miR-29TG mice (2 months) stained with H&E. Epidermis (E), dermis (D), subcutis (SC) and subcutaneous muscle (SM) layers are highlighted. **(E)** Increased collagen deposition (Masson’s Trichrome staining) in the skin of miR-29TG mice. **(F)** MicroCT scan images of the femur of wildtype and miR-29TG mice (2 months). **(G-I)** Femoral bone volume and bone mineral density were quantified (wildtype, n = 3; miR-29TG, n = 3). Cortical **(G)** and trabecular **(I)** bone volume (BV) are shown as a fraction relative to total bone volume (TV). Data shown are mean ± s.e.m., *p* values were calculated using an unpaired t-test. **(J)** Senescence-associated β-gal staining of the kidneys of young (2 months) wildtype, young (2-month-old) miR-29TG, and old (2-year-old) wildtype animals.

Increased cellular senescence is often associated with aging, and one of the most well-recognized senescence marker is positive staining with senescence-associated beta-galactosidase (SA-β-gal)^28^. To determine whether miR-29TG mice exhibit increased senescence, we conducted the SA-β-gal assay on kidneys isolated from young (2 months) miR-29TG, their WT littermates, as well as old (2 years) WT mice. In contrast to young WT controls, the miR-29TG kidneys showed strongly positive SA-β-gal staining similar to what was observed in old WT kidneys (Figure 3J). Taken together, these findings show that overexpression of miR-29 is sufficient to induce multiple features of an aging-like phenotype in mice.

To define the pathways that are altered in the miR-29TG mice, we conducted RNA-seq analysis of liver tissue from young (2 months) WT and miR-29TG mice. Principal component analysis showed that the mice stratify clearly by genotype status, irrespective of sex (Figure supplement 4A). We found a large number of genes to be differentially expressed between miR-29TG mice and WT mice (1928 genes downregulated and 2076 genes upregulated, FDR < 0.1) (Figure supplement 4B). The downregulated genes are highly enriched in fatty acid metabolism (Figure 4A), while the upregulated genes are highly enriched in pathways related to inflammation, including neutrophil activation (Figure 4B).

**Figure 4.**
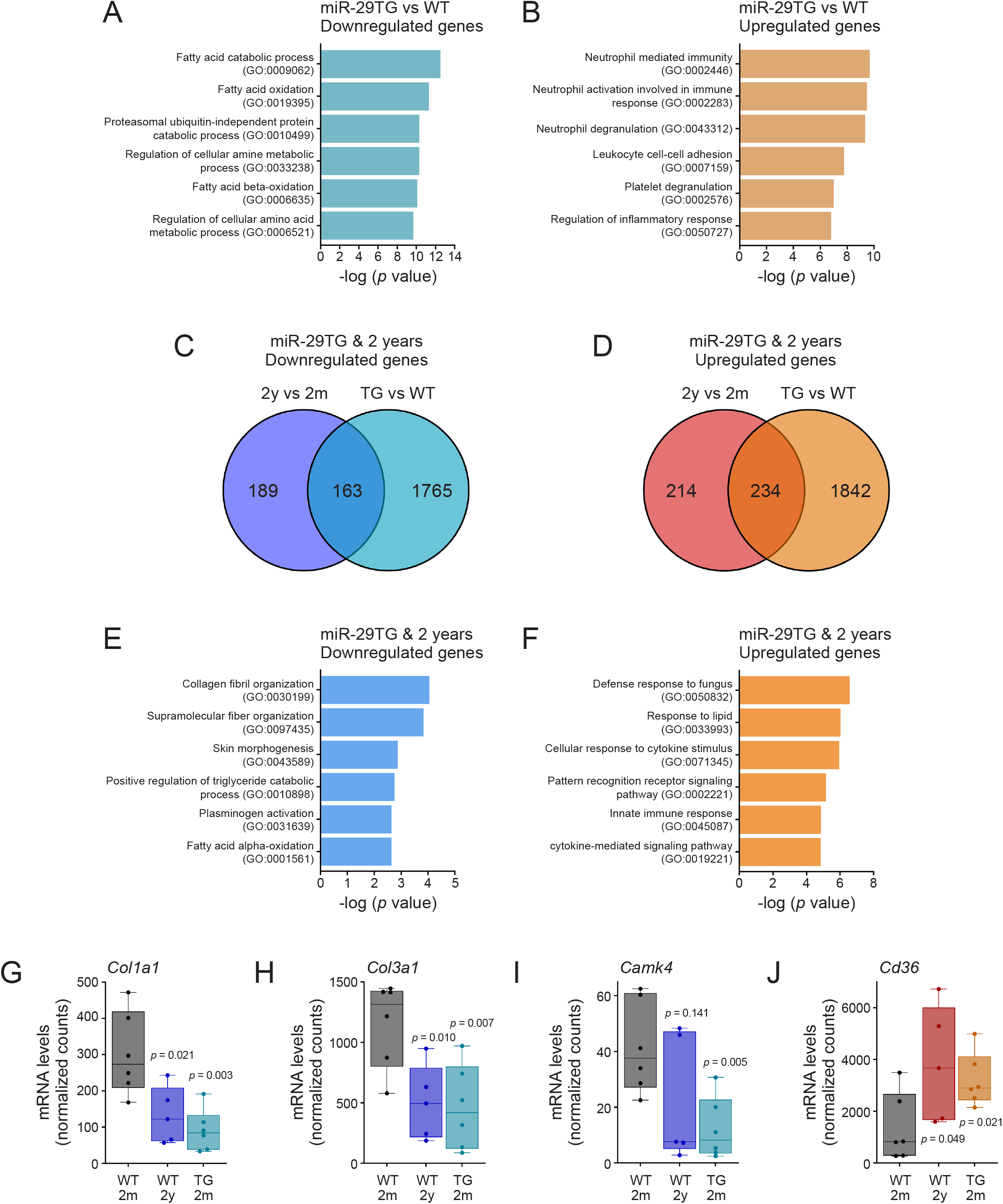
The liver of miR-29TG and 2 year-old wildtype mice share changes in key aging-related genes and pathways. **(A and B)** Pathway enrichment analysis of the genes that are significantly downregulated **(A)** and upregulated **(B)** in the liver of 2-month-old miR-29TG mice (n = 6; 5 males, 1 female) compared to 2-month-old wildtype mice (n = 6; 4 males, 2 females). The analysis was performed based on Gene Ontology (GO) Biological Process 2021. The top 6 processes are represented. **(C-J)** Comparison of the genes differentially expressed in the liver of 2-year-old wildtype mice and 2-month-old miR-29TG mice. **(C and D)** Number of genes downregulated **(C)** and upregulated **(D)** in common between the liver of 2-month-old (2m) miR-29TG and the liver of 2-year-old (2y) wildtype mice relative to 2-month-old (2m) wildtype mice. **(E and F)** Pathway enrichment analysis of shared downregulated genes **(E)** and upregulated genes **(F)** between the liver of 2-month-old miR-29TG and 2-year-old wildtype mice. Analysis was performed based on Gene Ontology (GO) Biological Process 2021. The top 6 processes are represented. **(G-J)** Expression of *Col1a1* **(G)**, *Col3a1* **(H)**, *Camk4* **(I)** and *Cd36* **(J)** mRNA in the liver of 2-month-old wildtype, 2-year-old wildtype and 2-month-old miR-29TG mice. The median, minimum and maximum are represented. The *p* values were calculated using an unpaired t test.

We next sought to identify the shared genes that are significantly altered in *both* young miR-29TG and old WT mice relative to young WT mice, as these genes may directly contribute to the aging phenotype. We detected 163 significantly downregulated (FDR < 0.1) and 234 significantly upregulated (FDR < 0.1) genes shared by young miR-29TG and old WT when compared to young WT mice (Figures 4C and 4D). Notably, we found that the shared set of up- and downregulated genes are significantly over-represented in many of the same pathways as the old WT mice (Figures 4E and 4F, Supplementary Tables 1, 2). Among the shared downregulated genes are collagen type 1 alpha 1 chain (*Col1a1*) and collagen type 3 alpha 1 chain (*Col3a1*) (Figures 4G and 4H), both of which are targets of miR-29^29,30^ and have been previously associated with aging^31-33^. Another shared downregulated gene is cAMP signaling calcium/calmodulin-dependent protein kinase IV (*Camk4*) (Figure 4I, Supplementary Table 1), a highly conserved predicted target of miR-29, which was previously found to be the most robustly downregulated gene in brains of aged mice, monkeys, and humans^34^.

A shared upregulated gene is *Cd36*, a membrane glycoprotein involved in fatty acid transport (Figure 4J, Supplementary Table 2). Previous studies have found *Cd36* to not only be upregulated in the hearts of aged mice but also to contribute to age-induced cardiomyopathy^35^. Interestingly, *Cd36* is also upregulated in response to senescence stimuli and is required for the senescence-associated secretory phenotype (SASP)^36^. Further, ectopic expression of *Cd36* alone is sufficient to induce many components of SASP ^36^ as well as induce senescence in young proliferating cells^37^. Interestingly, miR-29 has also been reported to induce cellular senescence *via* other pathways^38,39^. Thus, our study not only identifies genes that were previously known to be functionally associated with aging, but also reveals other genes (Supplementary Tables 1, 2) that could be important to drive aging-related phenotypes mediated by miR-29.

Taken together, our results highlight miR-29 as an important regulator of aging-related phenotypes. The concept that a single miRNA could regulate as complex a process as aging seems provocative, but individual miRNAs are well-recognized to have the ability to regulate very diverse cellular pathways^40,41^. While elevated levels of miR-29 have been previously reported to be associated with aging^1-10^, this is the first demonstration that miR-29 is sufficient to accelerate aging and drive aging-related phenotypes *in vivo*. Importantly, we also found reduction of miR-29 to extend lifespan in a progeria model.

Our knowledge about the pathways that are functionally important for regulating aging has come mostly from genetic studies in invertebrates and from mouse models of progeria. In mouse models of progeria, the mutations that cause aging are associated with either maintenance of nuclear architecture (*e*.*g*. Hutchinson-Gilford Progeria Syndrome) or genomic integrity (*e*.*g*. Werner’s syndrome, Cockayne syndrome)^42^. While many of the phenotypes seen in these progeroid models are also observed during normal aging, it is unclear whether the same genes drive the aging phenotype in both models. Our results suggest that miR-29 could be a candidate that regulates both pathological and physiological aging.

We propose that the pathways that are shared in multiple models of aging are likely to be more important for driving the aging phenotype. Our analysis of the genes shared between the miR-29TG mice and old mice highlights the downregulation of collagen synthesis (*e*.*g. Col1a1, Col3a1*), kinase signaling (*Camk4*), and the upregulation of *Cd36* as potential drives of the aging phenotypes. One mechanism by which *Camk4* could regulate aging is *via* its modulation of autophagy^43^, the reduction of which is a hallmark of aging^27^. This is consistent with the finding that the ortholog of miR-29 in *Caenorhabditis elegans*, miR-83, is elevated during aging and has been shown to reduce lifespan *via* inhibition of autophagy^44,45^. Pathway analysis also identified an upregulation of immune response in these models, a finding consistent with the well-recognized increase in inflammation in age-related disease and aging^27,46-50^. Interestingly, we found three components (*C6, C8A, C8B*) of the complement system’s Membrane Attack Complex (MAC) among the shared downregulated genes. Few studies have examined how the complement system changes with aging^51^. Our finding that multiple components of MAC are downregulated with aging suggests that this could be functionally linked with aging.

The upstream mechanisms by which miR-29 is elevated with age in mammals are not well defined. Recent work shows that Foxa2, a known transcriptional activator of miR-29^52^, exhibits increased chromatin occupancy in liver tissue of aged mice as well as of *Zmpste24*^*-/-*^ mice^53^. It remains unknown whether Foxa2 is the main driver of miR-29 during aging or whether other factors are involved. It is also unclear which tissues or cell types within a particular tissue are most relevant for the miR-29-mediated aging phenotype. Higher-resolution single-cell studies during aging as well as future experiments focusing on elevating miR-29 expression in specific tissues or cell types will help identify the specific context in which elevated miR-29 drives the aging-related phenotypes.

## Acknowledgements

The authors thank Dr. Peter J Havel (UC Davis) for sharing monkey tissue samples. We also thank Dr. Greg Hannon for sending us the mouse embryonic stem cells overexpressing miR-29b that we used to generate the miR-29TG mice. We appreciate the help of Mervi Eeva, Ying Li, and Bentley Midkiff at the UNC Translational Pathology Laboratory (TPL) for expert technical assistance and Dr. Dale Cowley for assistance in generating the miR-29 transgenic mice. We also thank Janice Weaver and Carolyn Suitt at the UNC Animal Histopathology and the Center for Gastrointestinal Biology and Disease (CGIBD), respectively, as well as Dr. Andrew Troy for assistance in isolating muscle tissue. The UNC TPL is supported in part by grants from the National Cancer Institute (P30CA016080) and the UNC University Cancer Research Fund. We thank the Small Animal Imaging Facility at the UNC Biomedical Imaging Research Center for providing the microCT imaging service. This work was supported by grants from the European Union (ERC-Advanced Grant, 742067), Ministerio de Ciencia e Innovación (Spain) (SAF2017-87655-R, PDI2020-118394RB-100), American Diabetes Association Pathway award 1-16-ACE-47 to PS, and NIH grants R01DK105965 to PS and NIH grants GM118331 and AG055304 to MD. XMC was supported by FPU fellowship (Ministerio de Educación, Spain).

## Supplementary Materials

Materials and Methods

Figures S1-S4

Tables S1-S2

Author contributions

## Materials and Methods

### Animals used in this study

Monkey liver samples were provided by the Havel lab at University of California-Davis. Tissues were collected from adult male rhesus macaques (age range 4 – 20.3 years), which were maintained at the California National Primate Research Center and provided a standard commercial nonhuman primate diet (5047; LabDiet, St. Louis, MO). All animal procedures were performed with the approval and authorization of the Institutional Animal Care and Use Committee (IACUC).

The *Lmna*^*G609G*^ knock-in and *Zmpste24*^*-/-*^mice were generated by the López-Otín lab. The *Lmna*^*G609G*^ knock-in mice carry a point mutation in the *Lmna* gene (1827C>T) that does not change the coding sequence (G609G) but activates a cryptic splicing donor site resulting in the generation of a truncated form of prelamin A (progerin). This point mutation in mice is equivalent to the mutation carried in Hutchibson-Gilford progeria syndrome patients^15^. The *Zmpste24*^*-/-*^ mice are deleted for the Zmpste24 gene, as described previously^17^. The *Lmna*^*G609G*^ and *Zmpste24*^*-/-*^ mice are both well-established models of progeria^54,55^.

The *miR-29a/b1*^*+/-*^ mice were also generated by the López-Otín lab as described^22^, from two embryonic stem (ES) cell lines carrying targeted deletions in the *miR-29a/b1* and *miR-29b2/c* clusters obtained from Dr. Haydn M. Prosser^56^. To generate the *miR-29a/b1*^*+/+*^ *Zmpste24*^*-/-*^ mice, we bred the *miR-29a/b1*^*+/-*^ with *Zmpste24*^*+/-*^ mice to obtain *miR-29a/b1*^*+/-*^ *Zmpste24*^*+/*-^ parental mice. By intercrossing *miR29a/b1*^*+/-*^ *Zmpste24*^*+/-*^, we obtained *miR-29a/b1*^*+/+*^ *Zmpste24*^*-/-*^ and *miR-29*^*+/-*^ *Zmpste24*^*-/-*^ at frequencies consistent with the expected Mendelian ratio. For the survival studies, mice were checked weekly. All animal experiments were approved by the Committee on Animal Experimentation of Universidad de Oviedo (PROAE 45/2018 and PROAE 46/2018).

### Generation of miR-29TG mice

Mouse embryonic stem cells overexpressing miR-29b used to generate the miR-29TG mice were obtained from Dr. Gregory Hannon (Cold Spring Harbor Laboratory). The parental ES cell line used to produce the miR-29-expressing clones was KH2 (C57BL/6 × 129/Sv). These cells are now commercially available at the NCI Mouse Repository (ES Cell Lines Catalog: M000108). The presence of the miR-29b transgene was confirmed by PCR with the following primers: 5’-CACCCTGAAAACTTTGCCCC-3’ and 5’-GCACCATTTGAAATCAGTGTTC-3’. The presence of the rtTA promoter was confirmed by PCR with the following primers: 5’-GGAGCGGGAGAAATGGATATG-3’, 5’-GCGAAGAGTTTGTCCTCAACC-3’, and 5’-AAAGTCGCTCTGAGTTGTTAT-3’. To induce the expression of miR-29, a mixture of doxycycline (2 mg/ml) and sucrose (50 mg/ml) was added to the drinking water of the mice starting on postnatal day 1. Mice were maintained in a 12 hrs light, 12 hrs dark cycle. The miR-29TG mice have been backcrossed to C57BL/6 mice and are maintained in the C57BL/6 background. All animal handling and protocols were carried out in accordance with established practices as described in the National Institutes of Health Guide for Care and Use of Laboratory Animals and as approved by the Animal Care and Use Committee of the University of North Carolina (UNC).

### Histology

For immunohistochemistry experiments, mice were anesthetized using isoflurane and transcardially perfused with 4% paraformaldehyde. The mice were then decapitated, and the various tissues were post-fixed in 4% paraformaldehyde overnight. Paraffin-embedded sections were used for hematoxylin-eosin (H/E), elastin, and Trichrome stains. Representative images are obtained from independent experiments, done in triplicate.

### cDNA synthesis and RT-qPCR analysis

RNA was extracted from various tissues using the Qiagen miRNeasy kit (catalog #217004). Mature miR-29b expression was assayed using TaqMan MicroRNA Assays (Applied Biosystems). Briefly, 10 ng of RNA was reverse transcribed using TaqMan miRNA reverse transcription kit (Applied Biosystems) and specific RT primers for miR-29b, U6 RNA, or snoRNA202 (Applied Biosystems). cDNA was amplified using a TaqMan Universal PCR Master Mix (Applied Biosystems). Relative quantification was carried out using the delta-delta Ct method. Sample variability was corrected by normalizing to U6 RNA (for WT and miR-29TG samples) or snoRNA202 (for WT and Lmna^G609G^ samples) levels.

### Computed tomography imaging

Computed tomography (CT) images were acquired using a custom-built Charybdis II scanner with a carbon nanotube field emission x-ray source. The source was operated at 50 kVp, with a 1.5 mA anode current and 50 ms exposure. 220 projections were acquired and reconstructed in COBRA (Exxim Computing Corporation, Pleasanton, CA). The image voxel size was 75.7 μm isotropic. For analysis of osteoporosis, samples were secured in a vertical orientation and scanned with a Scanco μCT40 ^57^ (Scanco Medical AG, Switzerland). Samples were scanned using the following parameters: 70 KvP, 114 μA, 8 W, 250 projections/slice, integration time 250 msec, and 12 μm voxel size. Standard midshaft and trabecular bone analyses were conducted using the Scanco Medical software. First, a region of interest was selected using the contouring tool. The cortical midshaft analysis consisted of 50 slices (600 μm) measured to be midway between the ends of the bone. For the trabecular analysis, 50 slices (600 μm) were taken approximately 200 μm from the end of the bone to avoid the growth plate. Regions of interest were drawn to include only trabecular bone in the analysis.

### Senescence-associated β-gal staining

Senescence-associated β-gal staining was performed with the Cellular Senescence Assay Kit according to the manufacturer’s instructions (Millipore).

### Flow cytometry analysis

Two-month-old WT and miR-29TG mice were euthanized and bone marrow, spleen and thymus cells were harvested following the standard procedure as previously described^58^. Specifically, single cell suspension of bone marrow cells was prepared by flushing the femurs and tibias using 3 ml ice-cold staining medium (Ca^2+^-and Mg^2+^-free Hank’s balanced salt solution, HBSS (Gibco) supplemented with 10 mM EDTA (Corning) and 2% heat-inactivated bovine serum (Gibco)) and filtered through 40 μm nylon mesh (Sefar). Thymocytes and splenocytes were isolated by crushing the whole thymus or spleen between two microscope slides followed by pipetting and filtering through nylon mesh. The number of viable cells was determined by diluting the cell with Turk Blood Diluting Fluid (Ricca Chemical) and manually counting using hemocytometer under microscope. To identify hematopoietic stem and progenitor cells by flow cytometry, bone marrow and spleen cells were stained with antibodies against c-kit (2B8, APC-Efluor780), Sca-1 (E13-161.7, PE-Cy7), CD48 (HM48-1, APC), CD150 (TC15-12F12.2, PE), CD16/32 (93, Alexa Fluor 700), as well as biotin conjugated antibodies against the following lineage markers: CD2 (RM2-5), CD3e (145-2c11), CD4 (GK1.5), CD5 (53-7.3), CD8a (53-6.7), B220 (RA3-6B2); Mac-1 (M1/70), Gr-1 (RB6-8C5), and Ter119 (TER-119), followed by staining with PE-CF594 conjugated streptavidin (BD Biosciences). To identify different T cell fractions in the thymus, total thymocytes were stained with ckit (APC-Efluor 780), CD4 (PE), CD8a (APC), CD44 (IM7, Alexa Fluor 700), CD25 (3C7, PerCP-Cy5.5) and biotin conjugated antibodies against the following lineage markers: B220, Mac-1, Gr-1, Ter119. All antibody staining were carried out on ice for 30 minutes. All primary and secondary antibodies were purchased from eBiosciences and Biolegend unless otherwise indicated. Flow cytometry analysis of stained cells was performed on a customized LSR II 7-laser, 17-color flow cytometer (Becton-Dickinson), and recorded data was analyzed in BD FACSDiva 8.0.1 and FlowJo 10.0.8 software.

### RNA library preparation and sequencing analysis

Liver was excised from P60 WT, P60 miR-29TG, and 2-year-old mice, and total RNA was isolated using the miRNeasy kit (Qiagen). RNA libraries were prepared using Illumina TruSeq (polyA+) and sequencing was performed on the Illumina HiSeq 2500 platform at the UNC High Through Sequencing Core Facility, yielding an average of ∼71 reads across samples. Sequencing reads were mapped to the GENCODE mouse transcriptome (vM10) using STAR (v2.4.2a) and transcripts were quantified using SALMON (v0.6.0). Across the samples, ∼80% of the reads were mappable. Transcript counts were normalized and differentially expressed genes (using FDR < 0.1) were identified using DESeq2 (v1.16.1). Raw sequencing data are available through GEO accession ID GSE107763.

### miRhub analysis

First, we used the seed-based target prediction algorithm TargetScanS 5.2^59^ to determine for each miRNA the number of predicted conserved targets among the genes in our gene sets. Each predicted miRNA-gene interaction was assigned a score based on the strength of the seed match, the level of conservation of the target site, the number of predicted target sites, and the clustering of target sites within that gene’s 3′ UTR. Finally, for each miRNA, the final targeting score was calculated by summing the scores across all genes and dividing by the number of genes. We repeated this procedure 10,000 times, with a new set of randomly selected mouse genes each time, in order to generate a background distribution of the predicted targeting scores for each miRNA (genes and corresponding 3′ UTR sequences were downloaded from http://www.targetscan.org). These score distributions were then used to calculate an empirical p-value of the targeting score for each miRNA in our gene set. To account for differences in the average 3′ UTR length between the genes of interest and the randomly selected genes in each simulation, the targeting score was normalized by 3′ UTR length.

### Pathway enriched analysis

Significantly differentially expressed genes (DEG) were defined as those with FDR < 0.1. Up- and down-regulated DEG lists were analyzed for enrichment in biological pathways using the online tool Enrichr^60^, which shows results from all available pathway databases. All pathway enrichment results shown in this manuscript are from the use of the Enrichr tool.

### Statistics

Statistical analyses were performed using GraphPad Prism 9 software. Data are expressed as mean ± s.e.m. in bar plots or median, minimum, and maximum in box and whiskers plots. The Student’s *t*-test (unpaired, two-tailed) for parametric data was used for analysis of two groups unless stated otherwise. No statistical methods were used to predetermine sample sizes, but our sample sizes are consistent with those reported in previous publications.

## Author Contributions

A.N. and V.S. conducted most of the experiments with help as described here. Y-H.H, M.K., C.L.K., J.B.G., and P.S. assisted in analyzing the RNAseq data. E.H., S.H., J.K., V.P., C.F., M.B., S.K., and N.S. assisted in conducting experiments that characterized the miR-29TG mice. X.M.C. and C.L.O. generated the miR-29-deficient mice and conducted the experiments involving the *Lmna*^*G609G*^ and *Zmpste24*^*-/-*^/miR-29-deficient mice. A.N., V.S., and M.D. outlined the project. M.D. and P.S. supervised the project, and E.H., A.N., V.S., P.S, and M.D. produced the final version of the manuscript.

## Supplementary Figure Legends

**Figure supplement 1.**
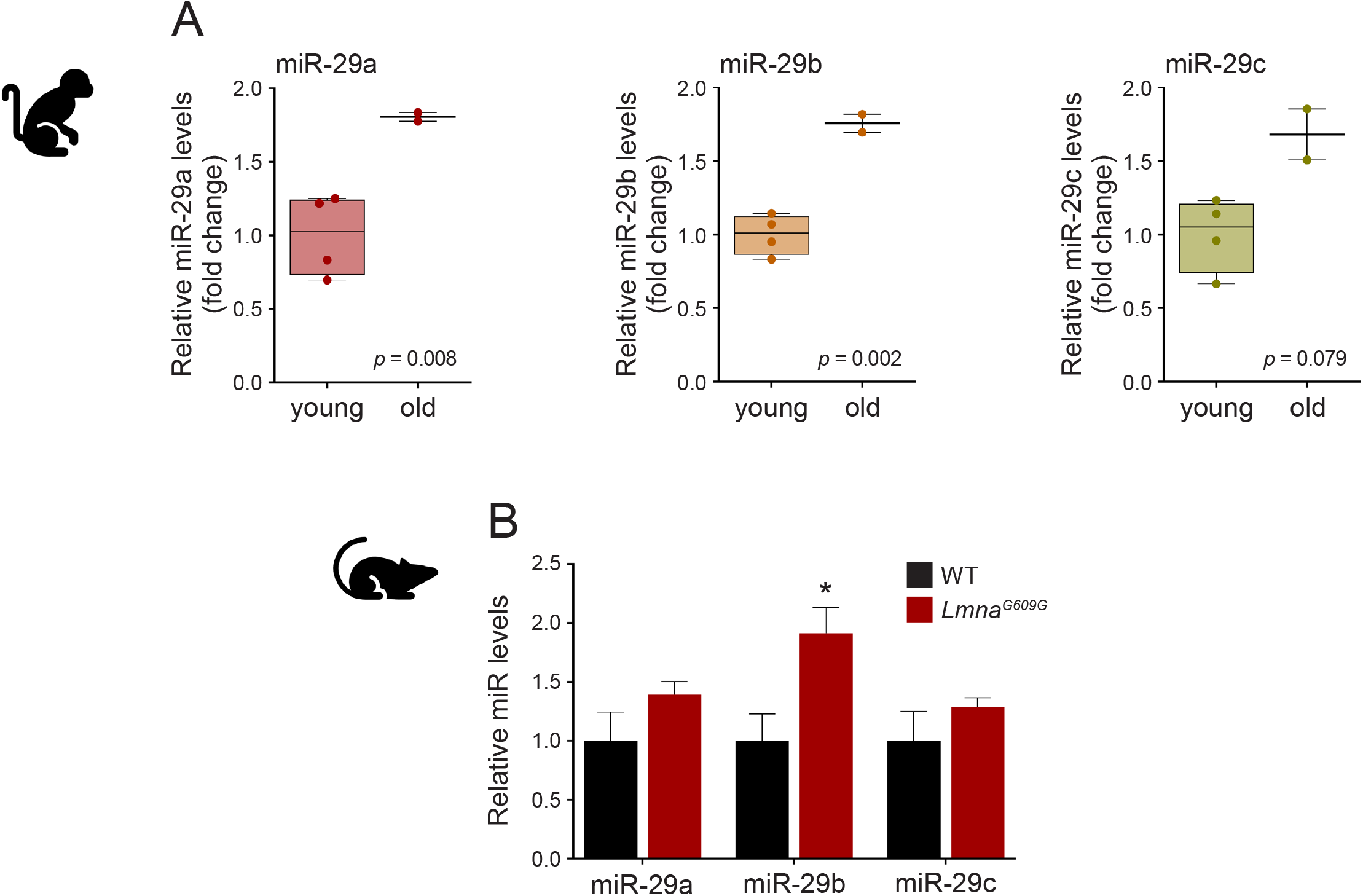
miR-29 expression increases with age in Rhesus macaque and in the *Lmna*^*G609G*^ mouse model of progeria. **(A)** Quantitative PCR analysis of miR-29 family members in the liver of young (4-8 year-old, n = 4) and old (14-20 year-old, n = 2) male Rhesus macaque. The median, minimum and maximum are represented. The *p*-value was calculated using an unpaired Welch’s t-test. **(B)** Relative levels of miR-29 family members in the heart of *Lmna*^*G609G*^ mice (n = 3). Data shown are mean ± s.e.m.

**Figure supplement 2.**
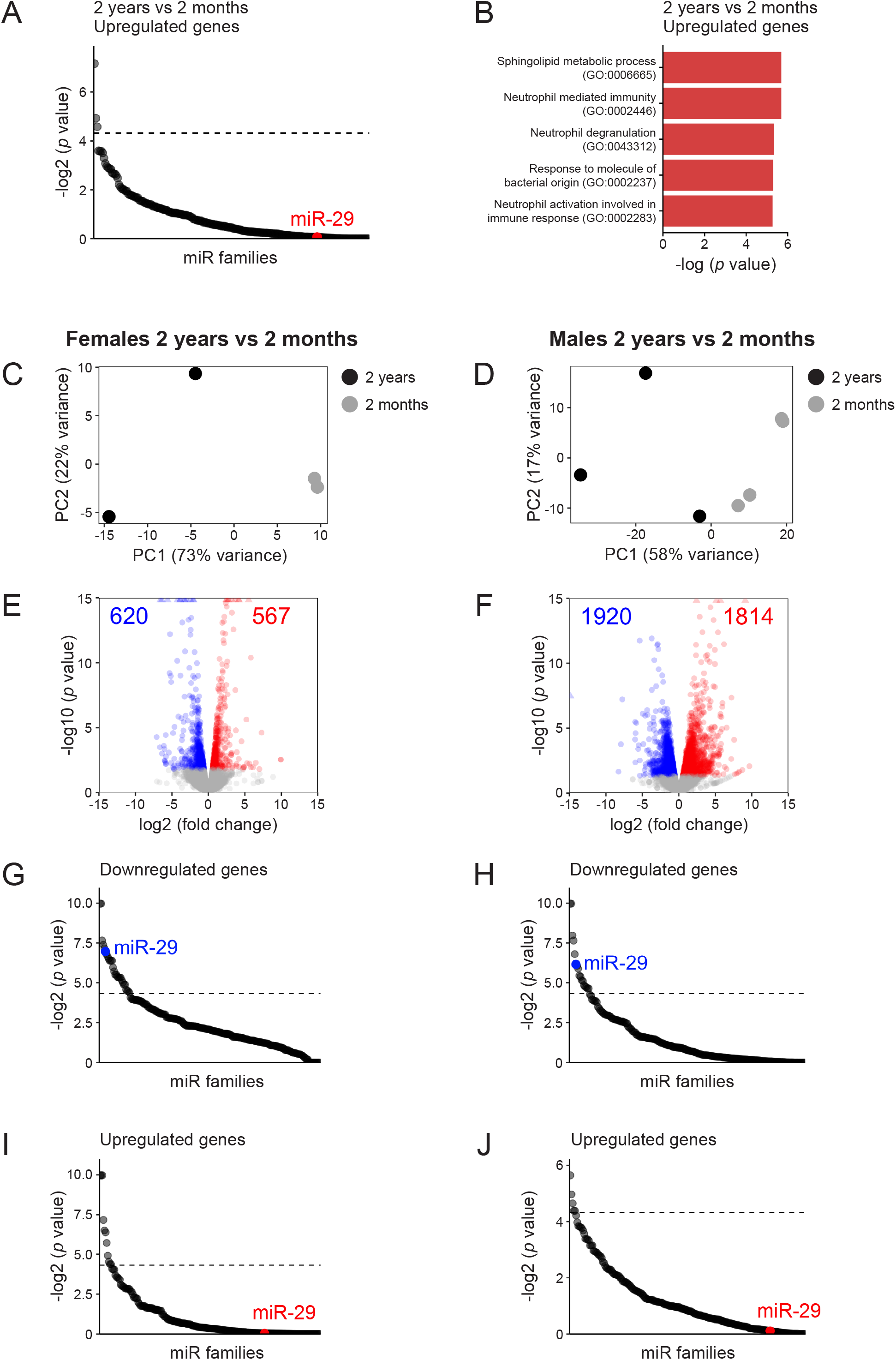
RNA-seq transcriptome analysis of the liver from old (2-year-old) and young (2-month-old) mice. **(A)** miRHub analysis of the downregulated genes in the liver of 2-year-old mice (n = 5; 3 males, 2 females) compared to the liver of 2-month-old mice (n = 5; 3 males, 2 females). Dashed line corresponds to *p* = 0.05. **(B)** Pathway enrichment analysis of the genes that are significantly upregulated in old compared to young mouse liver. The analysis was performed based on Gene Ontology (GO) Biological Process 2021. The top 5 processes are represented. **(C-J)** Sex-specific analysis of RNA-seq transcriptome profiles of liver from old (2-year-old, females, n = 2; males, n = 3) and young (2-month-old, females, n = 2; males, n = 3) mice. PCA plots for females **(C)** and males **(D)**. Volcano plot for females **(E)** and males **(F)**. The significantly upregulated and downregulated genes are represented in red and blue respectively. Filtering criteria: *p* < 0.05, adjusted *p* < 0.2 (Wald test; DESeq2). miRHub analysis of the downregulated genes in the liver of females **(G)** and males **(H)**. miRHub analysis of the upregulated genes in the liver of females **(I)** and males **(J)**. The predicted target sites of the miR-29 family are highlighted in blue for the downregulated genes **(G and H)** and in red for the upregulated genes **(I and J)**.

**Figure supplement 3.**
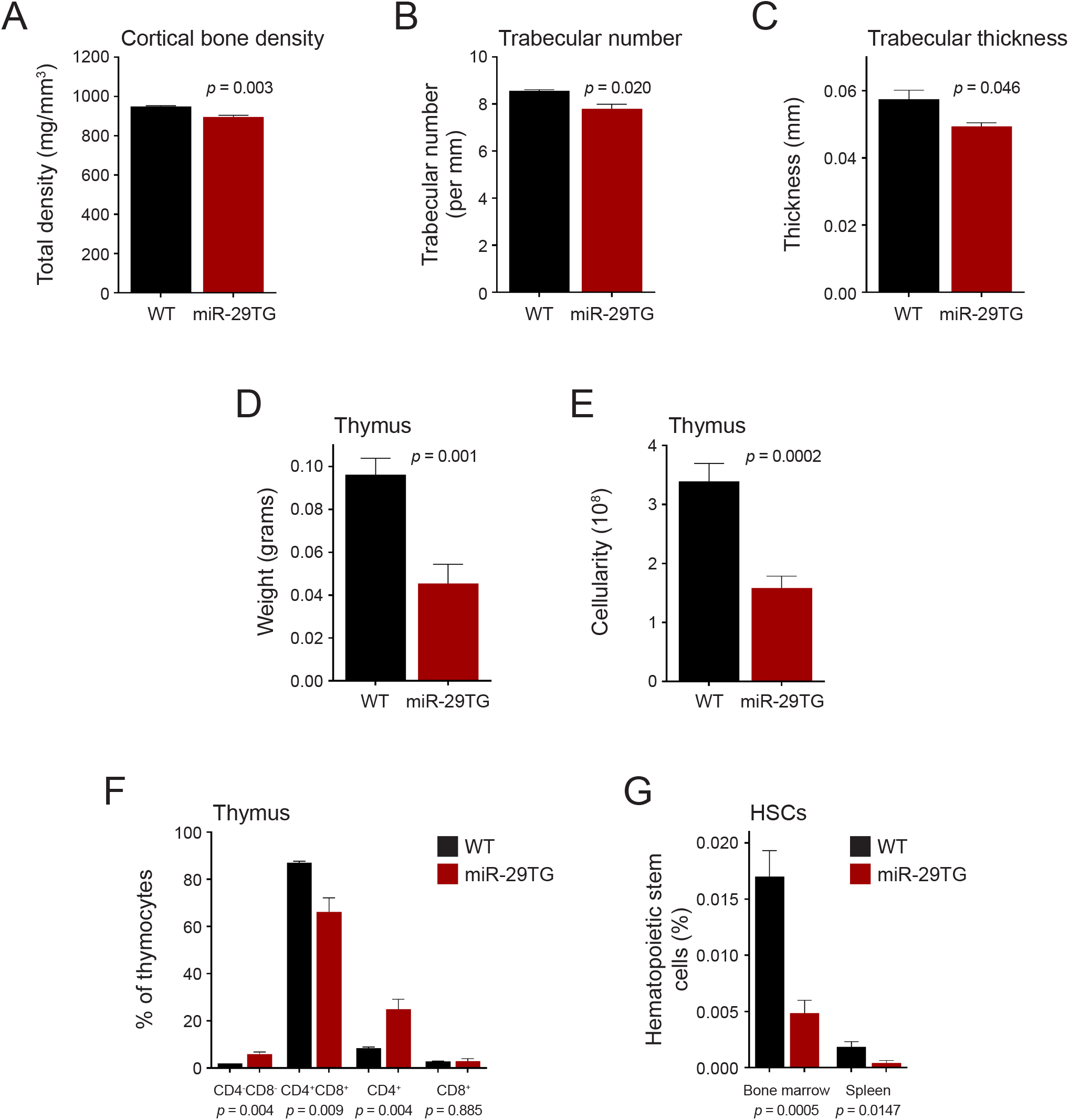
miR-29TG mice exhibit signs of osteoporosis and hematopoietic defects. **(A-C)** Quantification of microCT scan images of the femur of 2-month-old wildtype and miR-29TG mice (wildtype, n = 3; miR-29TG, n = 3). **(A)** Cortical bone density, **(B)** trabecular number and **(C)** trabecular thickness. **(D-F)** Analysis of thymus from 2-month-old wildtype (n = 7) and miR-29TG (n = 7) mice. **(D)** Thymus weights, **(E)** cellularity and **(F)** thymocyte lineages in WT and miR-29TG mice. **(G)** Bone marrow and spleen hematopoietic stem cell frequency in WT and miR-29TG mice. Data shown are mean ± s.e.m., *p* values were calculated using an unpaired t-test.

**Figure supplement 4.**
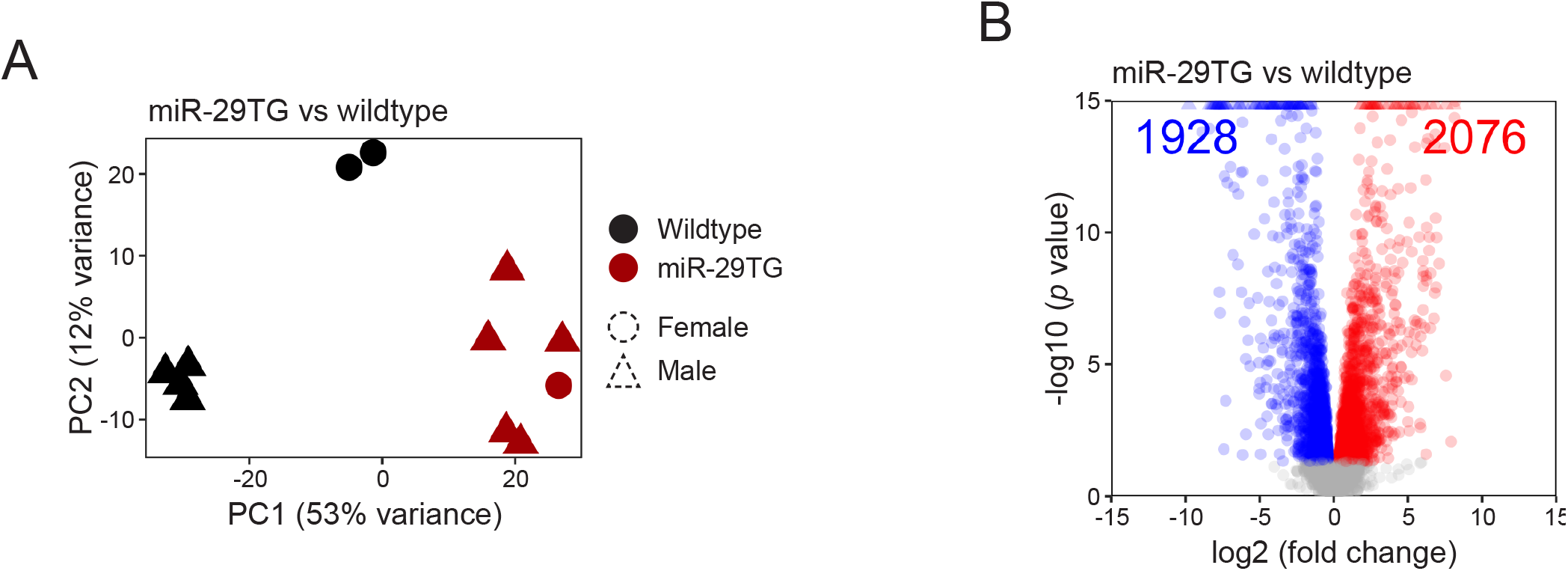
RNA-seq transcriptome analysis of the liver from young (2-month-old) miR-29TG and wildtype mice. **(A)** PCA plot of RNA-seq transcriptome profiles of the liver from 2-months-old miR-29TG (n = 6; 5 males, 1 female) and wildtype (n = 6; 4 males, 2 females) mice. Color denotes genotypes, shape denotes sex. **(B)** Volcano plot showing significantly upregulated (red) and downregulated (blue) genes in the liver of 2-month-old miR-29TG compared to 2-month-old wildtype mice. Filtering criteria: *p* < 0.05, adjusted *p* < 0.2 (Wald test; DESeq2).

## References and Notes

1 Ugalde, A. P. et al. Aging and chronic DNA damage response activate a regulatory pathway involving miR-29 and p53. EMBO J. 30, 2219–2232 (2011).

2 Takahashi, M., Eda, A., Fukushima, T. & Hohjoh, H. Reduction of Type IV Collagen by Upregulated miR-29 in Normal Elderly Mouse and klotho-Deficient, Senescence-Model Mouse. PLoS ONE 7, e48974 (2012).

3 Dimmeler, S. & Nicotera, P. MicroRNAs in age-related diseases. EMBO Mol. Med. 5, 180–190 (2013).

4 Boon, R. A. et al. MicroRNA-34a regulates cardiac ageing and function. Nature 495, 107–110 (2013).

5 Somel, M. et al. MicroRNA, mRNA, and protein expression link development and aging in human and macaque brain. Genome Res. 20, 1207–1218 (2010).

6 Xu, S. et al. The p53/miRNAs/Ccna2 pathway serves as a novel regulator of cellular senescence: Complement of the canonical p53/p21 pathway. Aging Cell 18, e12918 (2019).

7 Baumgart, M. et al. Age-dependent regulation of tumor-related microRNAs in the brain of the annual fish Nothobranchius furzeri. Mech. Ageing Dev. 133, 226–233 (2012).

8 Fenn, A. M. et al. Increased micro-RNA 29b in the aged brain correlates with the reduction of insulin-like growth factor-1 and fractalkine ligand. Neurobiol. Aging 34, 2748–2758 (2013).

9 Kamran, F. et al. Evidence That Up-Regulation of MicroRNA-29 Contributes to Postnatal Body Growth Deceleration. Mol. Endocrinol. 29, 921–932 (2015).

10 Caravia, X. M. & López-Otín, C. Regulatory Roles of miRNAs in Aging. Adv. Exp. Med. Biol. 887, 213–230 (2015).

11 Heid, J. et al. Age-dependent increase of oxidative stress regulates microRNA-29 family preserving cardiac health. Sci. Rep. 7, 16839 (2017).

12 Caravia, X. M., Roiz-Valle, D., Morán-Álvarez, A. & López-Otín, C. Functional relevance of Functional miRNAs in premature aging Mech. Ageing Dev. 168, 10–19 (2017).

13 Bates, D. J. et al. MicroRNA regulation in Ames dwarf mouse liver may contribute to delayed aging. Aging Cell 9, 1–18 (2010).

14 Kriegel, A. J., Liu, Y., Fang, Y., Ding, X. & Liang, M. The miR-29 family: genomics, cell biology, and relevance to renal and cardiovascular injury. Physiol. Genomics 44, 237–244 (2012).

15 Osorio, F. G. et al. Splicing-Directed Therapy in a New Mouse Model of Human Accelerated Aging. Sci. Transl. Med. 3, 106ra107 (2011).

16 Bergo, M. O. et al. Zmpste24 deficiency in mice causes spontaneous bone fractures, muscle weakness, and a prelamin A processing defect. Proc. Natl. Acad. Sci. USA 99, 13049–13054 (2002).

17 Pendas, A. M. et al. Defective prelamin A processing and muscular and adipocyte alterations in Zmpste24 metalloproteinase-deficient mice. Nat. Genet. 31, 94–99 (2002).

18 Pollex, R. L. & Hegele, R. A. Hutchinson–Gilford progeria syndrome. Clin. Genet. 66, 375–381 (2004).

19 Liu, B. et al. Genomic instability in laminopathy-based premature aging. Nat. Med. 11, 780–785 (2005).

20 Varela, I. et al. Accelerated ageing in mice deficient in Zmpste24 protease is linked to p53 signalling activation. Nature 437, 564–568 (2005).

21 Swahari, V. et al. MicroRNA-29 is an essential regulator of brain maturation through regulation of CH methylation. Cell Rep. 35, 108946 (2021).

22 Caravia, X. M. et al. The microRNA-29/PGC1α regulatory axis is critical for metabolic control of cardiac function. PLOS Biol. 16, e2006247 (2018).

23 Smith, K. M. et al. miR-29ab1 Deficiency Identifies a Negative Feedback Loop Controlling Th1 Bias That Is Dysregulated in Multiple Sclerosis. J. Immunol. 189, 1567–1576 (2012).

24 Rezzani, R., Nardo, L., Favero, G., Peroni, M. & Rodella, L. F. Thymus and aging: morphological, radiological, and functional overview. AGE 36, 313–351, doi:10.1007/s11357-013-9564-5 (2014).

25 Gui, J., Mustachio, L. M., Su, D. M. & Craig, R. W. Thymus Size and Age-related Thymic Involution: Early Programming, Sexual Dimorphism, Progenitors and Stroma. Aging Dis. 3, 280–290 (2012).

26 Fujino, T., Asada, S., Goyama, S. & Kitamura, T. Mechanisms involved in hematopoietic stem cell aging. Cell. Mol. Life Sci. 79, 473 (2022).

27 López-Otín, C., Blasco, M. A., Partridge, L., Serrano, M. & Kroemer, G. The Hallmarks of Aging. Cell 153, 1194–1217 (2013).

28 Imai, S. & Guarente, L. NAD+ and sirtuins in aging and disease. Trends Cell Biol. 24, 464–471 (2014).

29 Li, Z. et al. Biological functions of miR-29b contribute to positive regulation of osteoblast differentiation. J. Biol. Chem. 284, 15676–15684 (2009).

30 Chen, Y. et al. Cyclic stretch and compression forces alter microRNA-29 expression of human periodontal ligament cells. Gene 566, 13–17 (2015).

31 Lago, J. C. & Puzzi, M. B. The effect of aging in primary human dermal fibroblasts. PLoS ONE 14, e0219165 (2019).

32 Ohi, T., Uehara, Y., Takatsu, M., Watanabe, M. & Ono, T. Hypermethylation of CpGs in the promoter of the COL1A1 gene in the aged periodontal ligament. J. Dent. Res. 85, 245–250 (2006).

33 Bigot, N. et al. NF-κB accumulation associated with COL1A1 transactivators defects during chronological aging represses type I collagen expression through a -112/-61-bp region of the COL1A1 promoter in human skin fibroblasts. J. Invest. Dermatol. 132, 2360–2367 (2012).

34 Loerch, P. M. et al. Evolution of the Aging Brain Transcriptome and Synaptic Regulation. PLoS ONE 3, e3329 (2008).

35 Koonen, D. P. Y. et al. CD36 Expression Contributes to Age-Induced Cardiomyopathy in Mice. Circulation 116, 2139–2147 (2007).

36 Chong, M. et al. CD36 initiates the secretory phenotype during the establishment of cellular senescence. EMBO reports 19, e45274 (2018).

37 Saitou, M. et al. An evolutionary transcriptomics approach links CD36 to membrane remodeling in replicative senescence. Mol. Omics. 14, 237–246 (2018).

38 Martinez, I., Cazalla, D., Almstead, L. L., Steitz, J. A. & DiMaio, D. miR-29 and miR-30 regulate B-Myb expression during cellular senescence. Proc. Natl. Acad. Sci. 108, 522–527 (2011).

39 Hu, Z. et al. MicroRNA-29 induces cellular senescence in aging muscle through multiple signaling pathways. Aging (Albany NY) 6, 160–175 (2014).

40 Bartel, D. P. MicroRNAs: genomics, biogenesis, mechanism, and function. Cell 116, 281–297 (2004).

41 Bushati, N. & Cohen, S. M. microRNA functions. Annu. Rev. Cell Dev. Biol. 23, 175–205 (2007).

42 Kudlow, B. A., Kennedy, B. K. & Monnat, R. J. Werner and Hutchinson-Gilford progeria syndromes: mechanistic basis of human progeroid diseases. Nat. Rev. Mol. Cell Biol. 8, 394–404 (2007).

43 Zhang, X. et al. CaMKIV-Dependent Preservation of mTOR Expression Is Required for Autophagy during Lipopolysaccharide-Induced Inflammation and Acute Kidney Injury. J. Immunol. 193, 2405–2415 (2014).

44 Zhou, Y. et al. A secreted microRNA disrupts autophagy in distinct tissues of Caenorhabditis elegans upon ageing. Nat. Comm. 10, 4827 (2019).

45 Dzakah, E. E. et al. Loss of miR-83 extends lifespan and affects target gene expression in an age-dependent manner in Caenorhabditis elegans. J. Genet. Genomics 45, 651–662 (2018).

46 Neves, J. & Sousa-Victor, P. Regulation of inflammation as an anti-aging intervention. FEBS J. 287, 43–52 (2020).

47 McGeer, P. L. & McGeer, E. G. Inflammation and the degenerative diseases of aging. Ann. NY Acad. Sci. 1035, 104–116 (2004).

48 Chung, H. Y. et al. Molecular inflammation: Underpinnings of aging and age-related diseases. Ageing Res. Rev. 8, 18–30 (2009).

49 Ferrucci, L. & Fabbri, E. Inflammageing: chronic inflammation in ageing, cardiovascular disease, and frailty. Nat. Rev. Cardiol. 15, 505–522 (2018).

50 López-Otín, C. & Kroemer, G. Hallmarks of Health. Cell 184, 33–63 (2021).

51 Zheng, R. et al. The Complement System, Aging, and Aging-Related Diseases. Int. J. Mol. Sci. 23, 8689 (2022).

52 Kurtz, C. L. et al. MicroRNA-29 Fine-tunes the Expression of Key FOXA2-Activated Lipid Metabolism Genes and Is Dysregulated in Animal Models of Insulin Resistance and Diabetes. Diabetes 63, 3141–3148, doi:10.2337/db13-1015 (2014).

53 Whitton, H. et al. Changes at the nuclear lamina alter binding of pioneer factor Foxa2 in aged liver. Aging Cell 17, e12742 (2018).

54 Carrero, D., Soria-Valles, C. & López-Otín, C. Hallmarks of progeroid syndromes: lessons from mice and reprogrammed cells. Dis. Model Mech. 9, 719–735 (2016).

55 Folgueras, A. R., Freitas-Rodríguez, S., Velasco, G. & López-Otín, C. Mouse Models to Disentangle the Hallmarks of Human Aging. Circ. Res. 123, 905–924 (2018).

56 Prosser, H. M., Koike-Yusa, H., Cooper, J. D., Law, F. C. & Bradley, A. A resource of vectors and ES cells for targeted deletion of microRNAs in mice. Nat. Biotech. 29, 840 (2011).

57 Bouxsein, M. L. et al. Guidelines for assessment of bone microstructure in rodents using micro–computed tomography. J. Bone Mineral Res. 25, 1468–1486 (2010).

58 He, S. et al. Transient CDK4/6 inhibition protects hematopoietic stem cells from chemotherapy-induced exhaustion. Sci Transl Med 9, doi:10.1126/scitranslmed.aal3986 (2017).

59 Grimson, A. et al. MicroRNA Targeting Specificity in Mammals: Determinants beyond Seed Pairing. Mol. Cell 27, 91–105 (2007).

60 Kuleshov, M. V. et al. Enrichr: a comprehensive gene set enrichment analysis web server 2016 update. Nuc. Acid. Res. 44, W90–W97 (2016).

